# A psychophysically-tuned computational model of human primary visual cortex produces geometric optical illusions

**DOI:** 10.1101/2020.07.01.182329

**Authors:** Chrysa Retsa, Ana Hernando Ariza, Nathanael W. Noordanus, Lorenzo Ruffoni, Micah M. Murray, Benedetta Franceschiello

## Abstract

Geometrical optical illusion (GOIs) are mismatches between physical stimuli and perception. GOIs provide an access point to study the interplay between sensation and perception, yet there is scant quantitative investigation of the extent to which different GOIs rely on similar or distinct brain mechanisms. We addressed this knowledge gap. First, 30 healthy adults reported quantitatively their perceptual biases with three GOIs, whose physical properties parametrically varied on a trial-by-trial basis. Biases observed with one GOI were unrelated to those observed with another GOI, suggestive of (partially) distinct underlying mechanisms. Next, we used these psychophysical results to tune a computational model of primary visual cortex that combines parameters of orientation, selectivity, intra-cortical connectivity, and long-range interactions. We showed that similar biases could be generated *in-silico*, mirroring those observed in humans. Such results provide a roadmap whereby computational modelling, informed by human psychophysics, can reveal likely mechanistic underpinnings of perception.

## Introduction

The effortlessness of vision belies its mechanistic complexity as well as its perceptual fragility. Illusions are key phenomena to study vision. They provide a possibility to distinguish between sensation and perception. Illusions also have an ethological significance, as those organisms whose vision is capable of surmounting noise and ambiguity in visual scenes (e.g. camouflage) have a clear evolutionary advantage (Lesher, 1995). These phenomena also provide a more accurate depiction of the real world, playing a role in navigation (Lesher, 1995). Illusory figures present a solution to complex perceptual problems (Lesher, 1995; Rock & Anson, 1979). In this paper, we focus our attention on a subset of illusory phenomena called Geometrical-Optical Illusions (GOIs). They were first described in the 19th century by Johann Joseph Oppel (Oppel, 1855) and subsequently studied by other German psychologists such as Ewald Hering (Hering, 1861), Rudolph Hermann Lotze (Lotze, 1852) and Karl Zöllner (Zöllner, 1860).

GOIs were defined as situations where “there is a mismatch of geometrical properties between the item in the object space (physical source of the stimulus) and its associated percept” (Westheimer, 2008). Figure 1 (top row) presents some classical examples of GOIs: the Hering, the Zöllner and Poggendorff illusions. For example, in the Hering illusion (Figure 1, top row, A), the presence of a radial background induces a misperception of the two parallel lines, which appear as bowed towards the outside although they are straight. Optical illusions were first studied from the psychophysical point of view, leading to qualitative models that first identified the underlying mechanisms in terms of cognitive, retinal, and perceptual processes. Behavioral tests using these illusions have been performed during the 1960s-1980s (Beckett, 1989; Holt-Hansen, 1961; Oyama, 1975; Weintraub & Krantz, 1971), but there are both methodological limitations (e.g. hand-drawings were often used) and limited statistical analysis (i.e. most studies were purely observational and not quantitative). Therefore, the bias, i.e. *the difference between the geometrical nature of the stimuli and the perceived one* (amount of perceived curvature induced by the distance between original and corrected line, divergence and misalignment, respectively, for the GOIs depicted in Figure 1, top row), has not yet been investigated systematically. From the neurophysiological standpoint, several mechanisms have been proposed to contribute to illusory perceptual phenomena (Eagleman, 2001), such as lateral interactions between cells responsible for extracting basic features at the intracortical level, or feedback and feed-forward mechanisms across hierarchical levels of the visual pathway. Lateral inhibition refers to nearby neurons in the cortex spiking (near) simultaneously, due to competing features of the initial stimulus: the produced neuronal responses are themselves ambiguous in their coding. In the case of illusory contours, where geometrical objects such as lines or figures are perceived – although not physically present, feedback mechanisms have been postulated as – at least partly – responsible. Feedback modulations in the case of illusory perceptual phenomena allow for the incorporation of integration of information across longer distances, since higher-level cortices contain neurons with large receptive fields, and therefore the substantiation of Gestalt-like principles of perceptual grouping (Dura-Bernal, Wennekers, & Denham, 2011; Murray & Herrmann, 2013a). The specific contributions of these mechanisms to perceptual outcomes remain to be fully characterized.

**Figure 1.**
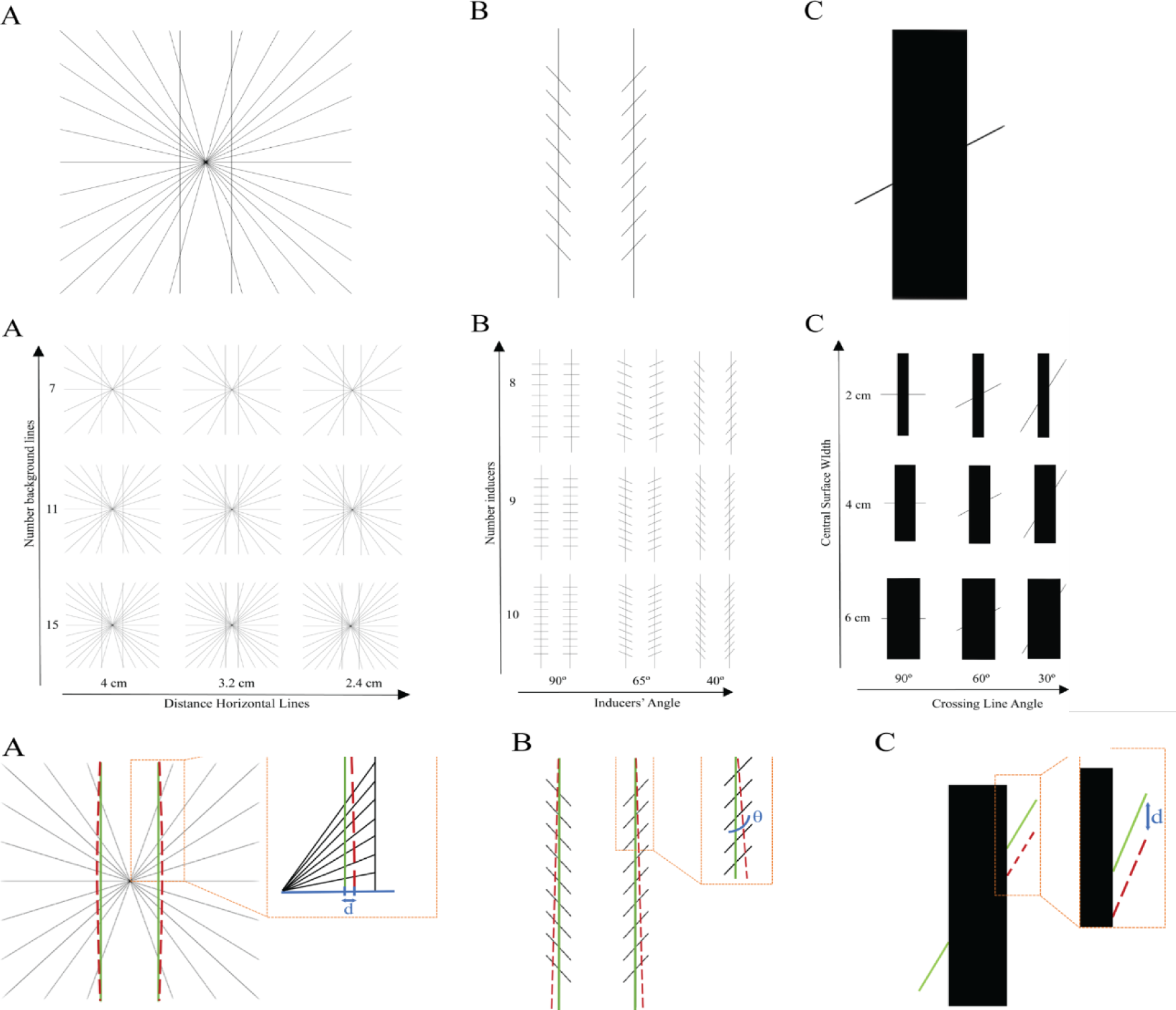
*(Top Row)* Examples of Geometrical-optical illusions. A) Hering illusion: two parallel vertical lines appear as curved due to the presence of a radial background. B) Zöllner illusion: two parallel vertical lines are perceived as non-parallel due to the presence of oblique inducers on the background. C) Poggendorff illusion: a perceived misalignment of the crossing oblique line is induced by the presence of a central surface. (Central Row) Visual stimuli used for the behavioral and the computational experiments. **A.** Hering illusion. The number of radial lines varies along the y-axis; the distance between the two vertical lines across the x-axis. **B.** Zöllner illusion. The number of inducer lines varies along the y-axis, the inducers’ angle across the x-axis. **C.** Poggendorff illusion. The width of the rectangle varies along the y-axis, the angle of the line segments across the x-axis. (Bottom Row) Bias calculation. Green line: geometrical real line. Red dotted line: corrected line. **A.** Hering illusion. The bias corresponds to the distance between the original line and the corrected line. **B.** Zöllner illusion. The bias refers to the angle (θ) created between the corrected and the parallel lines. **C.** Poggendorff illusion. The bias is the distance along the right rectangle edge between the corrected right line segment and the physically aligned right line segment.

Nonetheless, these psychophysical and neurophysiological observations have opened up the possibility of also investigating and computationally modeling the neural substrates of such illusory processing by means of the features (contour perception, brightness, depth) – determinants – of the stimuli (Lesher, 1995). Neural-based computational models of visual perception aim to provide mathematical descriptions of such neurophysiological features. A stable prediction of the misperception in GOIs could be explained by a model considering three main neurophysiological features (B. Franceschiello, Mashtakov, Citti, & Sarti, 2019; Benedetta Franceschiello, Sarti, & Citti, 2017): 1) orientation selectivity process performed by visual areas V1 and V2 (Hubel & Wiesel, 1962), 2) interference between receptive profiles of simple cells in these areas, and 3) long-range connectivity (Bosking, William H., 1997).

In terms of computational modelling, the orientation selectivity process is implemented by applying convolutional filters capable of extracting the most prominent orientation for each small portion of the visual field where the stimulus is represented. This interpretation holds in the case of possible interference between close-by receptive profiles. Many models have been proposed for this convolutional operation, among those the Derivative of Gaussians (DoG), introduced by Young (Young, 1987) and Daugman (Koenderink & van Doorn, 1990), Gabor filters, introduced by Daugman (Daugman, 1985) and Jones and Palmer (Jones & Palmer, 1987), and cake wavelets (Duits, Felsberg, Granlund, & Romeny, 2007). Gabor filters are particularly interesting as they have been proven to mimic accurately receptive profiles of simple cells in V1, as recorded neurophysiologically by De Angelis (De Angelis, Ohzawa, & Freeman, 1995). As for the long-range connectivity (or horizontal connectivity) of the primary visual areas, we refer to the mechanism “connecting” neurons spiking simultaneously while situated at relatively far distances (ca. 300-400*μm*) from one another. Long-range connections enable the formation of contours, surfaces, and more complex objects and they have been mathematically described in terms of mean field equations (Bertalmío et al., 2020; Bertalmío & Cowan, 2009; Bressloff, Cowan, Golubitsky, Thomas, & Wiener, 2001; Schuster & Wagner, 1990), of statistics of the visual inputs (Bednar, 2014; Simoncelli & Olshausen, 2001) or by a geometrical description of the visual areas (Ben-Shahar & Zucker, 2004; Citti & Sarti, 2006; Hoffman, 1980; Petitot, 2002). Among those computational models providing a description for GOIs, we find a statistical-based approach in (Fermüller & Malm, 2004) and a geometrical approach in (Ehm & Wackermann, 2012, 2016). The first models interpreting GOIs while having a neural-based approach are those in (Franceschiello, Benedetta, Sarti, Alessandro, Citti, 2017; Franceschiello et al., 2019). These provided a first *qualitative* reconstruction of the phenomena. By integrating psychophysical data from actual observers, we were therefore in a position to determine the contributions of the neural components of the *in-silico* simulations, providing for the first time a *quantitative* reconstruction of the phenomena.

The contributions of the present paper are threefold. Firstly, and to our knowledge for the first time, we introduce a standardized, quantitative measure of perceptual bias for each of the three GOIs: Hering, Zollner and Poggendorff illusions. Furthermore, these perceptual illusions are analyzed together for the first time, enabling us to determine to what extent biases are related and by extension the degree to which distinct mechanisms contribute to each illusion. Second, we establish the practice of tuning the computational models in (B. Franceschiello et al., 2019; Benedetta Franceschiello et al., 2017) using psychophysical data: computational models are strongly dependent on the choice of parameters. A part of them is only computationally driven, while another part represents neurophysiological correlates of the studied processes. Tuning the computational model means iteratively optimizing the parameters to achieve a replication of human behavior. The third contribution of this work is to derive neural-based conclusions from those neurophysiological-related parameters which permitted the computational model fitting, i.e. a quantitative reproduction of the phenomena. This allows us to finally reveal those mechanisms responsible for the perception of geometrical optical illusions.

## Materials and Methods

### Behavioral Experiment

#### Participants

Thirty healthy unpaid volunteers (19 females; aged 22– 39years; mean±SD=28.4 ±4.3years) provided informed consent to participate in the experiment. All procedures conformed to the 2013 update of the Declaration of Helsinki and were approved by the Cantonal Ethics Committee. None of the subjects had current or prior neurological or psychiatric illnesses. All participants had normal or corrected-to-normal vision.

#### Stimuli

The experiment included stimuli for the three different illusion tasks: Hering, Zöllner and Poggendorff. Visual stimuli were independently varied along 2 aspects of interest (i.e. our independent variables for each condition). For each illusion, nine sets of visual stimuli were presented as in different conditions, following a 3×3 factorial design.

#### Hering stimuli

The neutral configuration of Hering stimuli consists of two parallel vertical lines (foreground curves) and a symmetrical arrangement of radial background lines, which meet in the center of the image, halfway between the two foreground lines. The participants then usually report the two straight, parallel vertical lines as appearing curved (Hering, 1861). In the present experimental set-up, participants had to adjust the vertical lines in order to appear subjectively straight and parallel.

In our experiment, each stimulus is drawn in a rectangular region in the center of the screen with a gap above and below of 5% of the height of the screen (the height of the screen used was 20 cm), such that the vertical extent of the stimulus is the middle 90% of the height of the screen. The background of the stimulus consists of a number of lines disposed radially, see Figure 1, central row, (A). These lines are evenly spaced along the edges of the central square, as a rule always including the horizontal and diagonal lines and excluding the central vertical line (see Figure 1, central row, (A)). All stimuli consisted of black lines presented on a white background.

The nine different stimuli in the Hering condition were obtained by varying independently the number of radial lines and the distance between the two main vertical lines (Figure 1, central row, (A)). These two variables were chosen because they constitute the main components of the Hering illusion. Each stimulus included either 7, 11 or 15 radial lines (the corresponding values in the Psychopy code for stimuli presentation are 5, 7 and 9) and a distance between the two vertical lines of either 2.4 cm, 3.2 cm and 4 cm, (the corresponding values in the code are: 0.06, 0.08, 0.1). The values were fixed in order to have perceivable differences while experiencing the illusory effects. To recover the measure in cm, these values (0.12, 0.16 and 0.2) were multiplied by the height of the screen (20 cm).

The two foreground curves are not always drawn straight, but the curves appear smooth in all configurations. The foreground curves can veer towards or away from the center of the image. We define the offset to be the initial horizontal distance (varied for each trial) between one of the straight vertical line in the neutral configuration and the corresponding foreground curve (see Appendix A for further explanations). We varied the initial value of the offset by discrete steps (up to 4 steps in either direction). When the participant adjusts the offset parameter by pressing the up or down key, the value of the offset is adjusted by 1 step. We varied the initial value of the offset in order not to bias the participant towards a stable initial configuration. However, this parameter is not among our main experimental manipulations. The measure of this experiment is the final offset, which corresponds to the distance from the neutral configuration and represents what the participant perceives as straight/parallel, and it is measured in discrete steps. This is the perceived bias (Figure 1, bottom row, A).

#### Zöllner stimuli

The neutral configuration of Zöllner stimuli consists of two vertical lines each intersected by a number of segments (inducers), see Figure 1, central row, (B). Despite the fact that the vertical lines are parallel to each other, the participants perceive them as not parallel and instead report them to appear to converge and diverge from each other. In the present experimental set-up, participants had to adjust the angle of the vertical lines in order to appear subjectively parallel.

In our experimental setup, the inducers are parallel segments spread evenly over the length of the vertical lines. The inducers intersecting the left vertical line are consistently oriented from top left to bottom right and the inducers on the right line form their mirrored image. The length of the vertical lines is 80% of the height of the screen (16cm). The distance between them is 4 cm (0.2 of the screen height). The length of the inducers is 2cm (0.1 of the screen height).

The nine different stimuli in Zöllner were obtained by varying independently the two following parameters: the number of inducers and the angle of the inducers, Figure 1, central row, (B). In our example, only the angle is manipulated and the number of inducers, which could vary between 8, 9 and 10 inducers, is used as control. The angle can vary between 40°, 65° and 90° (from vertical). These latter values were fixed in order to have perceptual differences while experiencing the illusory effects. The participants manipulated the angle of the vertical lines (both lines were rotated together in a mirrored way). The offset step size is 0.1 degrees. The vertical lines in the Zöllner illusion were not always drawn parallel; sometimes they were tilted by a small angle, which is considered as the distance from the neutral configuration; it is what we refer to as offset and it varies in discrete steps. We varied the initial offset between −0.6 ° to 0.6 °. When the participant adjusts the offset parameter by pressing the up or down key, the value of the offset is adjusted by 1 step. The measure of this experiment is called final offset, which corresponds to the distance from the neutral configuration and represents what the participant perceives as parallel. This is their perceived bias (Figure 1, bottom row, (B).).

#### Poggendorff stimuli

The neutral configuration of Poggendorff stimuli consists of two collinear segments separated by a central surface. The participants usually report the two collinear segments as misaligned. In the present experiment, participants had to adjust the vertical position of the right segment in order for the two segments to be subjectively perceived as collinear.

In our experimental setup, the stimuli consist of a central black rectangle, with a fixed height of 16 cm and a width that varies from one presentation to another, and two black lines, one on the left side and one on the right. The nine different stimuli in Poggendorff were obtained by varying independently the width of the rectangle and the angle (slope) of the line segments, Figure 1, central row, (C). These two independent variables were chosen as they constitute the main components of the Poggendorff illusion. The width of the rectangle was varied between 2, 4 and 6 cm, and the angle of the line segments between 30°, 60° and 90° (from the vertical configuration). These values were chosen in order to have a significant difference in the perception of the illusory effects. The two segments always have the same slope. Both line segments extend to 4 cm horizontally away from the center of the screen. Therefore, their length depends on the width of the rectangle and on the slope. In the neutral position (zero offset), the right and left segments both align with the central point of the rectangle, as if to form a single line passing through that point. The left line is always fixed in this position. However, as in the other illusion tasks, there is an initial offset variable. Specifically, the right line can be offset by a variable amount ranging between −0.04 to 0.04 cm. When the participant adjusts the offset parameter by pressing the up or down key and moving the right line segment upwards or downwards, the value of the offset is adjusted by 1 step of 0.02 cm size. The measure of this experiment is the final offset, which is the vertical distance from the neutral configuration (alignment of the segments) and represents what the participant perceives as collinear. This is the perceived bias (Figure 1, bottom row, (C)).

#### Procedure and Experimental design

Participants sat in a darkened, sound-attenuated room (WhisperRoom MDI, 102126E) in front of a laptop screen (MacBook Pro end 2013, 13-inch, resolution: 2560 x 1600) that was presenting the different stimuli. The distance between participants’ eyes and the screen was kept constant at approximately 75 cm, for a vertical visual angle of 15°, which implies that the stimuli were projected in participants’ central vision. The experiment consisted of three different blocks, one for each illusion task. Participants performed each illusion task one after the other; the order of which was counterbalanced across participants. Participants took short breaks between the different tasks. Within each block, the nine different sets of stimuli were presented randomly to the participants. Each stimulus remained on the screen for as long as the participant took to respond. Participants then pressed the spacebar in order to proceed to the next trial. The order of trials was randomized for each participant. Short instructions were presented at the beginning of each of the tasks, followed by a couple of practice trials for the participant to get familiarized with the tasks. Stimulus delivery and behavioral response collection were programmed and controlled by PsychoPy v3.0 (Peirce et al., 2019). The code for the experiments is publicly available at the following link: https://gist.github.com/nat-n/879d06244a694d47c4911098aac60656.

Depending on the illusion, participants were instructed to adjust the target lines (green lines in Figure 1, bottom row) in order to i) either make them appear subjectively parallel (Hering and Zöllner) or ii) to appear collinear (Poggendorff).

To move the target lines, their task was to press either the up or down arrow keys on the keyboard. In the Hering illusion, the target lines were the two foreground curves (Figure 1, bottom row, (A)). These curves were bent outwards if the down button was pressed and inwards if the up key was pressed, according to the mathematical expression of the curve (see Appendix, section A). For the Zöllner illusion, the target lines were again the two vertical lines. The vertical lines shifted outwards if the up key was pressed and inwards if the down key was pressed (Figure 1, bottom row, (B)). In the Poggendorff illusion, the target line was the right line segment (Figure 1, bottom row, (C)). The right segment moved up if the up key was pressed and down if the down key was pressed.

#### Hering design

A 3 (number of radial lines: 7, 11 and 15) by 3 (distance between vertical lines: 2.4 cm, 3.2 cm and 4 cm) factorial design was adopted, resulting in 9 experimental conditions. Each of the nine conditions was presented 12 times, resulting in 108 trials in total. It should be noted that the initial value of the offset was not specified as an experimental factor. For each condition, we set several initial offset values: – 0.107 cm (1 time), −0.053 cm (3 times), 0 cm (4 times), 0.053 cm (3 times) and 0.107 cm (1 time), following a Gaussian distribution with mean equal to the condition we wanted to test most often. The bias here was defined as the horizontal distance between the original vertical line and the manipulated final curve, see Figure 1, bottom row, (A). An explanation that links the described “distance” and the curvature of the final curve is presented in the Appendix, (A). To calculate the distance, we summed the number of times the up and down arrow keys were pressed, giving it a positive value when the down key was pressed and a negative value when the up key was pressed. The resulting final offset, encoding the final position of the line according to the participant’s perception, was then multiplied by the offset step size, equal to 0.0267 cm in the Hering illusion (see the Stimuli section).

#### Zöllner design

A 3 (number of inducers: 8, 9 and 10) by 3 (angle of inducers: 40°, 65° and 90°) factorial design was adopted, resulting in 9 experimental conditions. Each of the nine conditions was presented 12 times, resulting in 108 trials in total. The initial value of the offset was not specified as an experimental factor. For each condition, we set several initial offset values: – 0.6° (1 time), −0.4° (3 times), 0° (4 times), 0.4° (3 times) and 0.6° (1 time), following a Gaussian distribution with mean equal to the condition we wanted to test most often. The bias referred to the angle created between the corrected and the parallel lines Figure 1, bottom row, (B). To calculate that angle, we summed the number of times the up and down arrow keys were pressed, giving it a positive value when the up key was pressed and a negative value when the down key was pressed. The resulting value (final offset) was then multiplied by the angle in degrees corresponding to how much the line moved each time we pressed an arrow, i.e. its offset step size equal to 0.1 °.

#### Poggendorff design

A 3 (width of the rectangle: 2, 4 and 6cm) by 3 (angle of the line segments: 30°, 60° and 90°) factorial design was adopted, resulting in 9 experimental conditions. Each of the nine conditions was presented 12 times, resulting in 108 trials in total. The initial value of the offset was not specified as an experimental factor. For each condition, we set several initial offset values: – 0.04 cm (1 time), −0.02 cm (3 times), 0 cm (4 times), 0.02 cm (3 times) and 0.04 (1 time), following a Gaussian distribution with mean equal to the condition we wanted to test most often. The bias for Poggendorff was calculated as the distance, computed along the right side of the rectangle, between the right line segment and the actual physically aligned segment, Figure 1, bottom row, (C). To calculate that distance, we summed the number of times the up and down arrow keys were pressed, giving it a positive value when the up key was pressed and a negative value when the down key was pressed. The resulting value (final offset) was then multiplied by the distance in cm that the line moved each time we pressed an arrow (offset step size equal to 0.02 cm)

#### Behavioral analysis

IBM SPSS Statistics 25 (IBM Corp. Released 2017. IBM SPSS Statistics for Windows, Version 25.0. Armonk, NY: IBM Corp.) was used for data analysis. First, we tested whether the recorded data were normally distributed using a Shapiro-Wilk test. All data were normally distributed. Hence, a 3 × 3 repeated measures ANOVA was used for each illusion task in order to analyze the effects of our experimental manipulations on the participants’ calculated biases. The factors that were used were: the number of radial lines (7, 11, 15) and the distance between the vertical lines (2.4 cm, 3.2 cm, 4 cm) for Hering, the number of inducers (8, 9, 10) and angle of inducers (40°, 65°, 90°) for Zöllner and the width of the rectangle (2 cm, 4 cm, 6 cm) and the angle of the line segments (30°, 60°, 90°) for Poggendorff.

### Computational experiment

#### Theoretical formulation of the model

The cortically inspired computational models introduced in (B. Franceschiello et al., 2019; Benedetta Franceschiello et al., 2017) have proven to be effective in reproducing the qualitative responses of the primary visual cortices in response to geometrical optical illusions. The model, tested in (B. Franceschiello et al., 2019; Benedetta Franceschiello et al., 2017) from a purely image processing perspective, takes into account as a first step the orientation selection mechanisms discovered by Hubel and Wiesel (Hubel & Wiesel, 1962) and modelled computationally by Daugman (Daugman, 1985) and Jones and Palmer (Jones & Palmer, 1987). For a full description of the model, we refer readers to (B. Franceschiello et al., 2019; Benedetta Franceschiello et al., 2017). Here, we briefly summarize the approach. Gabor filters have the following formulation in their expression independent from time:

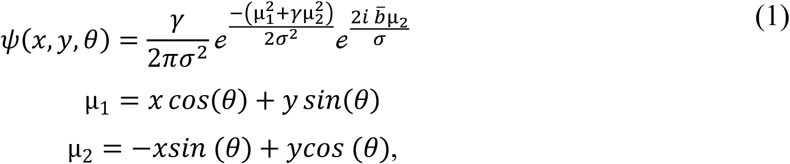

where (*x, y*) ∈ ℝ^2^ represents the global coordinates on the initial image; 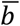, *σ* and *γ* are extra parameters determining the shape and size of the filter. *ψ*(*x, y, θ*) is a complex-valued function, therefore we can identify a real part and imaginary part. The real part is an even function with respect to (*x, y*):

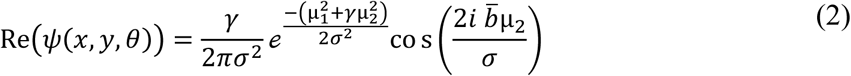

While the imaginary one an odd function with respect to (*x, y*):

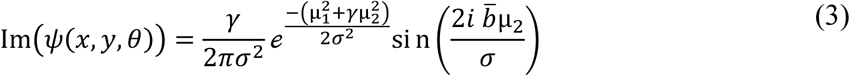

Odd and even parts of Gabor filters describe different types of receptive field profiles of simple cells observed in V1. Odd receptive field profiles are able to detect contours and therefore are used to detect the presence of a surface in an image, such as the central surface presented in the Poggendorff illusion. In this case, *θ* ∈ [−*π, π*). Even receptive field profiles are sensitive to line orientation, and for this reason *θ* ∈ [0, *π*).

As shown by Citti and Sarti (Citti & Sarti, 2006), the organization of simple cells in V1 responds to a group law that allows to retrieve (*x, y, θ*) position of simple cells receptive fields in the hypercolumnar structure of V1. (*x, y, θ*) is retrieved by means of a rotation and a translation of a mother cell placed at the origin of our space, ℝ^2^×S^1^, where ℝ^2^ encodes the spatial features while S^1^ the orientation selection. If we identify the retinal plane with the ℝ^2^-plane, then the spike frequencies *O* (x, y, θ) of the neurons activating at the global coordinates (x, y) are modeled as the convolution between the visual stimulus *I*: *D* ⊂ ℝ^2^ → ℝ^+^ with the set of Gabor filters. The expression for this output becomes the following:

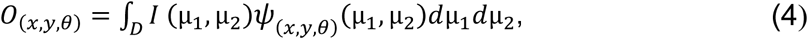

where 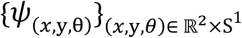 is the set of Gabor filters, *D* ⊂ ℝ^2^. By means of this mechanism, we mimic the orientation selectivity process, i.e. the short-range or intracortical connectivity, as described in the experimental works of Hubel and Wiesel (Hubel & Wiesel, 1962). In the Hering and Zöllner illusions, the maximum output among all possible instances of orientations is given by *E*(*x, y, θ*) *=* ‖*O*(*x, y, θ*)‖, i.e. the Energy, or Complex module of the output presented in Equation (4). This output is specific for line detection, therefore *θ* ∈ [0, *π*). In the Poggendorff illusion, the output has the following expression:

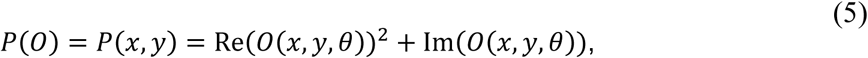

where again *O* (*x, y*, θ) is the output of simple cells defined in Equation (4). This formulation facilitates the detection of both the presence of lines, thanks to the contribution of the Real part in Equation (2) and the presence and polarity of contours thanks to the term in Equation (3). Therefore *θ* ∈ [−*π, π*). *P*(*x, y*) is then normalized and shifted to positive values, obtaining the following external cost *C*:

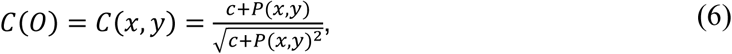

where *c* is a suitable positive constant. Both the Energy *E*(*x, y, θ*) and the external cost *C(O)* simulate the intracortical connectivity of V1/V2.

Once every simple cell in the features space ℝ^2^×S^1^ has been assigned with an output by means of the simulated intra-cortical connectivity (*E*(*x, y, θ*) and *C(O)*), the next step is to explain how long-range horizontal connections, i.e. connections between cells of approximately *the same orientation* belonging to different hypercolumns (and by extension responding to different regions of the visual field) is taken into account by the model. Citti and Sarti (Citti & Sarti, 2006) proposed to endow the ℝ^2^×S^1^ features space with a sub-Riemannian metric, which allows us to weigh differently the direction of propagation of the long range connectivity of V1 and V2. In (B. Franceschiello et al., 2019; Benedetta Franceschiello et al., 2017) the model presented in (Citti & Sarti, 2006) is extended to take into account how the long-range connectivity is polarized by the output of simple cells. As the nature of the illusions is different, two different approaches are proposed, as detailed in (B. Franceschiello et al., 2019; Benedetta Franceschiello et al., 2017).

#### Hering and Zöllner illusions

A connectivity metric tensor is defined by combining a long-range connectivity metric in ℝ^2^×S^1^ with the output *E*(*x, y, θ*), for every orientation instance of the hypercolumn, as follows:

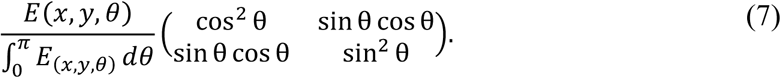

The second part of the expression represents a 2-dimensional connectivity metric tensor along the directions {∂_x_, ∂_y_}, in the roto-translation group coordinate system. Summing up (integrating) the contribution along the hypercolumn we obtain a 2-dimensional deformation tensor with principal eigenvector along the orientation 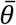, corresponding to the maximum output of the energy within the hypercolumn:

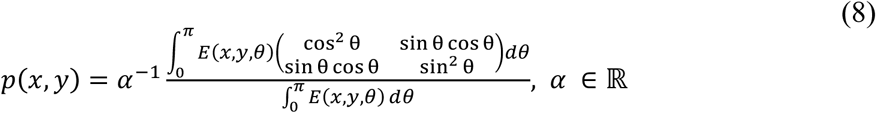

By plugging in the components of the inverse matrix of *p*(*x, y*) inside the following equation:

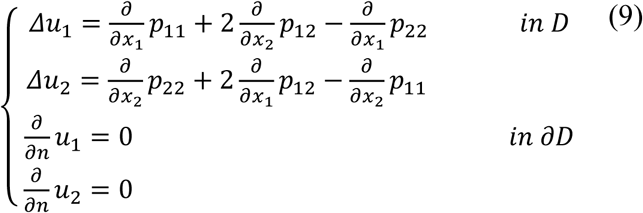

it is possible as detailed in (Benedetta Franceschiello et al., 2017) to recover the displacement field *ū = (u*_*1*_, *u*_*2*_), see Figure 2 (by solving (9)). The differential equation introduced in the formula before approximates the connectivity patterns between different hyper-columns as some diffusive propagation of single cells responses, complementing the intra-cortical connectivity patterns.

**Figure 2.**
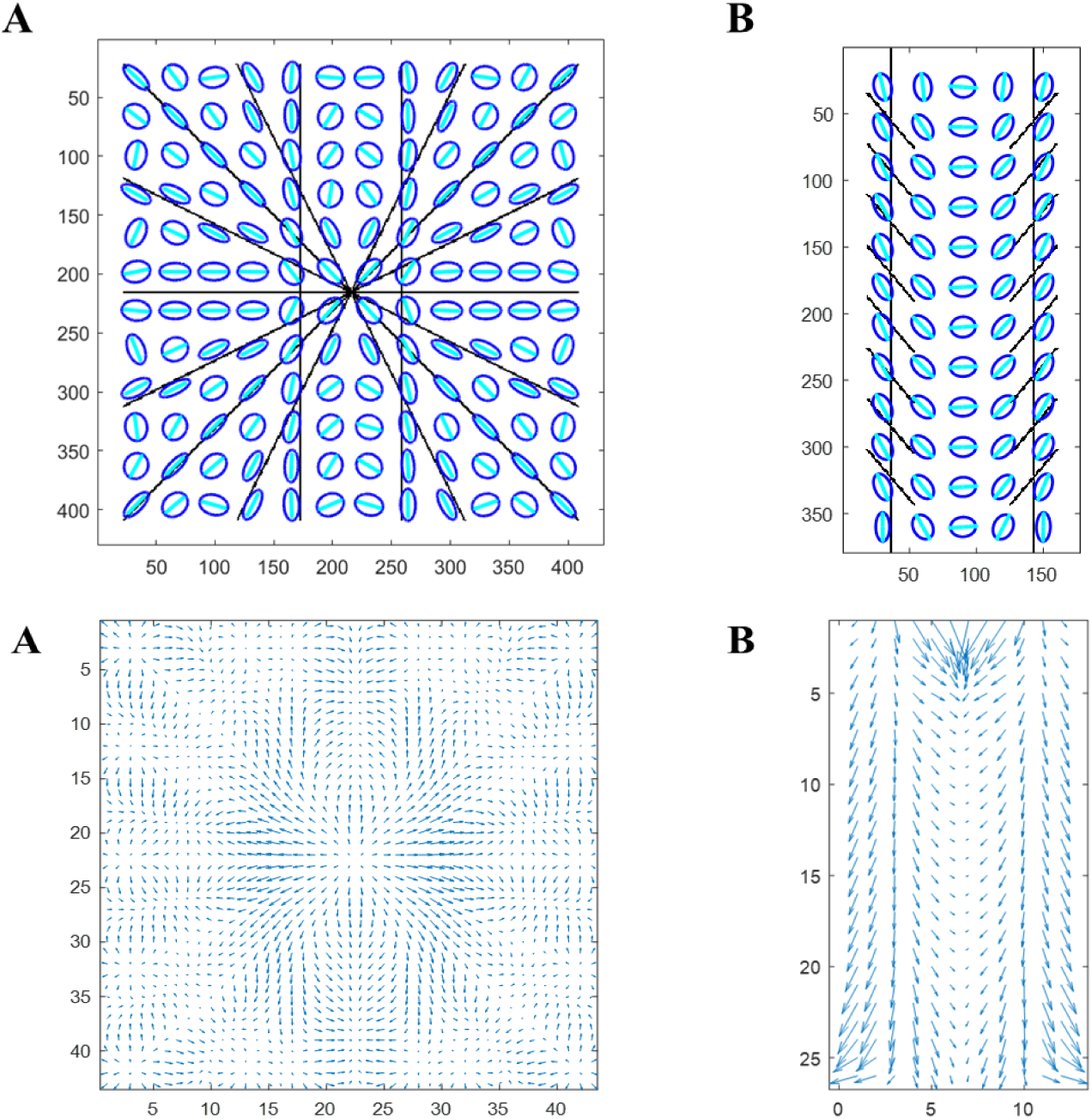
(Top row) Representation of p. In blue, representation of a tensor field. In cyan, the eigenvector corresponding to the principal eigenvalue. **A.** Example of one stimulus of the Hering illusion (7 background lines and 4cm distance between vertical lines). **B.** Example of one stimulus of the Zöllner illusion (10 inducers and 65° for the inducer’s angle). (Bottom row) Computed displacement field *ū*. **A.** Example for one of the stimuli of the Hering illusion (7 background lines and 4cm distance between vertical lines). **B.** Example of one stimulus of the Zöllner illusion (10 inducers and 65° for the inducer’s angle).

#### Poggendorff illusion

The basic idea presented in (B. Franceschiello et al., 2019) is to provide a natural environment for this type of illusion by means of geometrical elements and to model the perceptual curves as length minimizers of a cortical metric. Starting from the sub-Riemannian metric in (Citti & Sarti, 2006), we weigh the long-range connectivity considering the intra-columnar response *P*(*O*), see Equation (5), of simple cells in V1/V2. The polarized metric becomes:

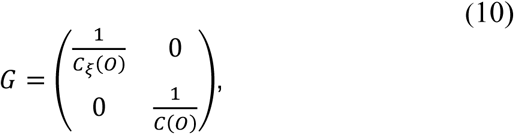

where *C*_ξ_(*O*) = ξ^2^*C*(*O*), *C*(*O*) is defined in Equation (6). The top left term weights the ℝ^2^ component of the space while the bottom right the rotational S^1^ component. ξ > 0 is a real parameter, therefore allowing to modulate the anisotropy between the retinal (on ℝ^2^) and the hypercolumnar (S^1^) components. In the case of the Poggendorff illusion, the reconstructed perceived misalignment is described by the minimizing geodesic between two a priori known sets, obtained by fixing the point (x,y, θ) on the left side of the surface and looking for another point (x_1_, y_1,_ θ) on the right side of the surface, where we fix everything except for y_1_. The illusory curves will be minimal with respect to the metric induced by the geometry of the primary visual cortex, i.e. the deformation curves arise as geodesics of the metric *G*, Equation (10).

Computation of sub-Riemannian distances in general is a very difficult problem (Montgomery, 2006). By introducing a Riemannian approximation of the metric, the problem becomes solvable and geodesics (perceptual curves) arise as solutions of the Riemannian Eikonal equation (Mirebeau, 2018; Sanguinetti et al., 2015).

#### Numerical Implementation

For the implementation of the mathematical models described above, we use MATLAB ver. R2019a. The initial images used for the behavioral test were extracted and resized manually, to optimize the computation time dependent on the size of the initial image. For the three illusions, the original stimuli were images of size 2880×1800 pixels, first cropped along the horizontal and vertical directions to obtain a square of size 1800×1800 pixels (Hering), a rectangle of 592×1267 pixels (Zöllner) and a rectangle of 770×1540 pixels (Poggendorff). The three images were then scaled by a factor 4.18, 3.34 and 3.85, respectively, obtaining final images with sizes 430×430 pixels (Hering), 177×379 pixels (Zöllner) and 200×400 pixels (Poggendorff). The resized images corresponding to the stimuli were convolved with a set of Gabor filters (N=36 orientations for the Hering and Zöllner, N = 32 for the orientation of lines extraction in the Poggendorff illusion and N=62 for the surface contour orientation extraction).

The number N of orientations stated above was computed as equally spaced steps within the two intervals, the latter being *θ* ∈ [0, *π*) (for the Hering, the Zöllner and the Poggendorff) and *θ* ∈ [−*π, π*) (for the Poggendorff), depending on the features extracted (i.e. lines or contours as detailed above).

The implementation of the Gabor filters is the one presented in (Petkov & Subramanian, 2007), setting the required parameter *λ =* 0. According to Daugman (Daugman, 1985), Gabor filters simulate the receptive field profiles of simple cells in V1/V2 responsible for the orientation tuning. The size of the Gabor patch, i.e. the dimensions of the filter windows in pixels, was varied across illusions by tuning *σ*. The implementation of Gabor filters in Matlab is available in http://www.cs.rug.nl/~imaging/spatiotemporal_Gabor_function/GaborKernel3d.m (Petkov & Subramanian, 2007). The Gabor filters’ parameters were fixed for all presentations of the same illusion: *γ =* 0.5; 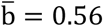; *σ =* 7.84 for the Hering; *γ =* 0.5; 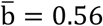; *σ =* 11.2 for the Zollner; *γ =* 1; 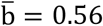; *σ =* 3 for the orientation of lines extraction in the Poggendorff; and *γ =* 1; 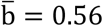 *=* 0.56; *σ =* 7 for the orientation of contours extraction in the Poggendorff. These parameters were fixed based on previous computational studies in accord with a qualitative reconstruction of the percept (B. Franceschiello et al., 2019; Benedetta Franceschiello et al., 2017). In the Hering and Zöllner illusion a constant *c =* 0.037, 0.03 respectively scales the displacement vector fields *ū* = (x_1_, x_2_) once applied to the initial image to reconstruct the percept. In the Poggendorff experiment, the parameter ξ – which modulates the anisotropy between the two direction, ξΔx_1_ = Δθ, where Δx_1_, Δθ are the discretization steps along x_1_ and θ – is chosen based on the entry angle for the transversal line and the width of the central surface. The geodesics are computed through the Sub-Riemannian Fast-Marching, as implemented in (Sanguinetti et al., 2015). The parameter *ϵ* indicates the Riemannian approximation and in our experiments is set equal 0.1. Table 1 shows the values of ξ for the presented experiments.

**Table 1.** Chosen values for *ξ*

| Central Width | Crossing Line Angle | $\xi$ |
| --- | --- | --- |
| 2 cm | 30° | 30 |
|  | 60° | 13 |
|  | 90° | 10 |
| 4 cm | 30° | 30 |
|  | 60° | 13 |
|  | 90° | 10 |
| 6 cm | 30° | 30 |
|  | 60° | 20 |
|  | 90° | 17 |

#### Bias calculation in the computational results

The computational model, once tuned, was fitted with the outcome of the behavioral results according to the variable of interest of the three detailed geometrical optical illusions. As our goal is to compare the results obtained in the behavioral experiments with the resulting biases of the computational protocol, the definitions of bias are the same as those given in the behavioral bias paragraph. However, the way the bias is extracted in the computational case is clearly different, but it has been standardized in order to reduce any variability induced by a manual extraction of the measures.

#### Computational bias Hering illusion

To recall the definition, the bias is the horizontal distance between the straight vertical line of the neutral configuration and the curve computed by the model. To calculate it in a standardized way, we extracted the distance between point 1 and 2, see Figure 3 (A) for reference, which corresponds to the value in pixels of the displacement vector field *ū = (u*_*1*_, *u*_*2*_) computed at point 1, multiplied by the constant *c* =0.037, see *Numerical Explanation section*.

**Figure 3.**
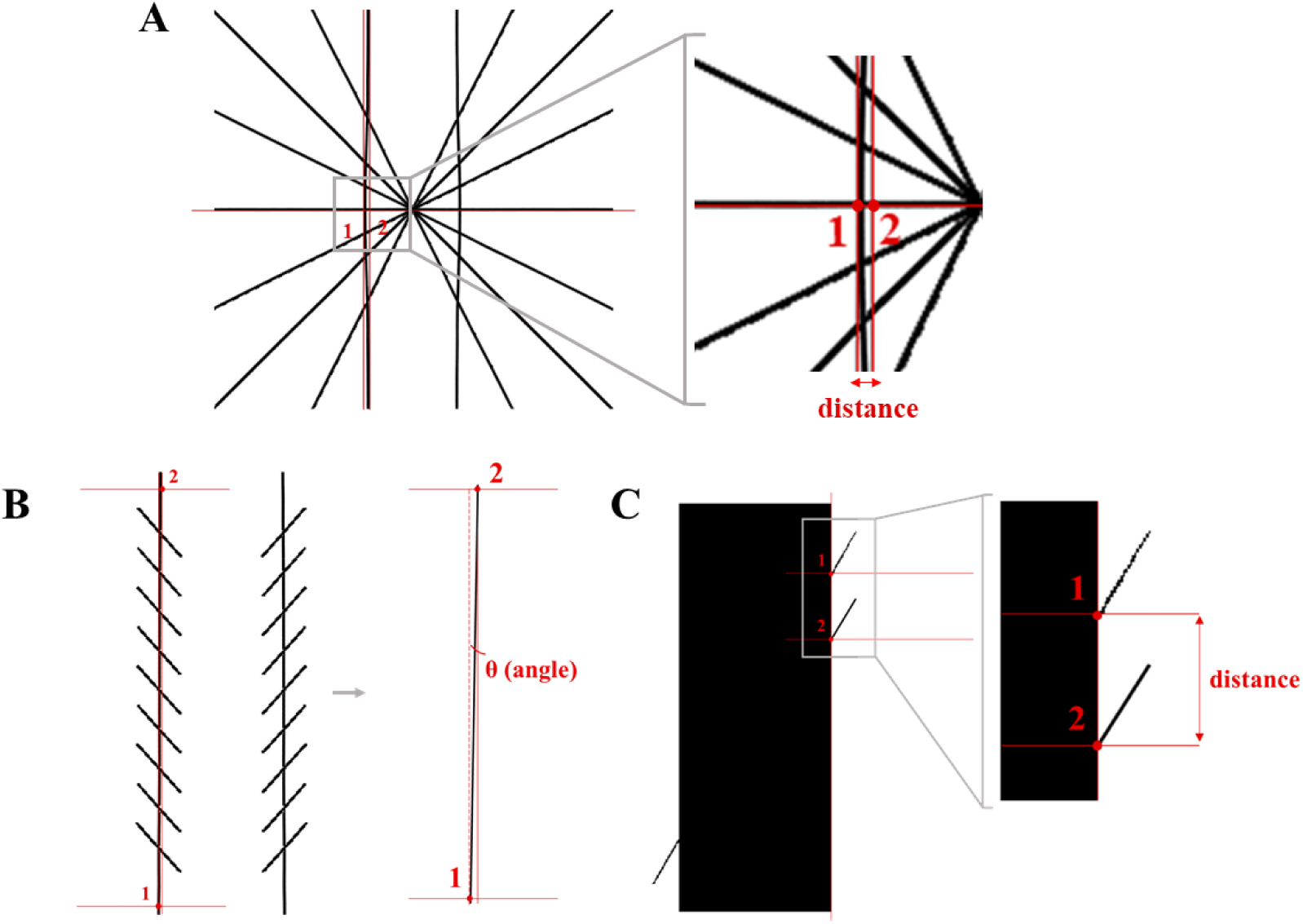
Bias calculation for the computational modelling. 1 and 2 indicate the two reference points for the bias calculation. **A.** Hering illusion. The bias corresponds to the distance between the original vertical line position and the tangent in the point of the maximal curvature. **B.** Zöllner illusion. The bias refers to the angle rotation (θ) of the modelled line. **C.** Poggendorff illusion. The bias is the distance along the right side of the central surface between the initial position of the right segment line and the modelled one.

#### Computational bias Zöllner illusion

The bias refers to the angle created between the modelled vertical line and the original parallel one. To calculate the angle, we choose two points in the final image: the first one on the bottom of the left of the left perceptual reconstructed line (*x*_1_, *y*_1_), and the second on the top of the same line (*x*_2_, *y*_2_), see Figure 3 (B). The rotation angle is computed as atan 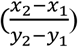. The result is then converted from radians to degrees.

#### Computational bias Poggendorff illusion

The bias for Poggendorff is again calculated as the vertical distance between the reconstructed segment corresponding to the perceptual alignment and the neutral configuration (in which the left and right segments are aligned). The computational bias then results as the difference in pixels between the original y-coordinate of the physically aligned segment and the y-coordinate of the segment obtained through our algorithm (Figure 3 (C)).

### Tuning the computational model through behavioral results

As described in the numerical implementation section, the computational models used to simulate the perceptual behavior of the visual cortex that generates the perception of geometrical optical illusions depend on a series of parameters. Such parameters do have an interpretation from the neurophysiological standpoint and therefore impact the fitting of the computational curves with the behavioral ones. By *tuning the computational models*, we refer to the practice of iteratively modifying these parameters in a fashion that would lead to optimally fit the computational biases with the behavioral ones. This practice is particularly advantageous in the case of neural-based models, like in our case, i.e. those models that depend on parameters characterizing low-level visual processes, including simple-cell feature extraction and long-range connectivity.

### Comparison between computational and behavioral biases

The computational results were then compared with the behavioral ones to tune the model and evaluate its prediction. Different sets of initial parameters for the size of receptive fields (encoded by the *σ* of the Gabor filter) were tried before choosing those described in the *Numerical implementation* subsection. The final value of *σ* was chosen as the instance maximizing the fitting between computational and behavioral curves (minimizing the error), and remained fixed across different conditions of the same illusion. The comparison between computational and behavioral biases of the Zöllner illusions was straightforward, as the compared variables are both in degrees. To ensure the same comparison between the distances extracted in the Hering and Poggendorff illusions, the bias is expressed in percentages; in the Hering illusion as 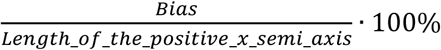; in the Poggendorff illusion is computed as 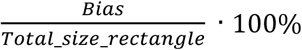. In this way, the behavioral measures extracted in cm become comparable with the computational measures in pixels.

## Results

### Behavioral results

Results of the psychophysical experiments are shown in Figure 4. Biases were calculated as detailed in Sect. 2.1.4. The average size of the illusion (bias) for each task was submitted to a 3 × 3 repeated measures ANOVA (see 2.1.4), as the distribution of the data was Gaussian. Reported F and p values reflect Greenhouse-Geisser correction for non-sphericity when necessary.

**Figure 4.**
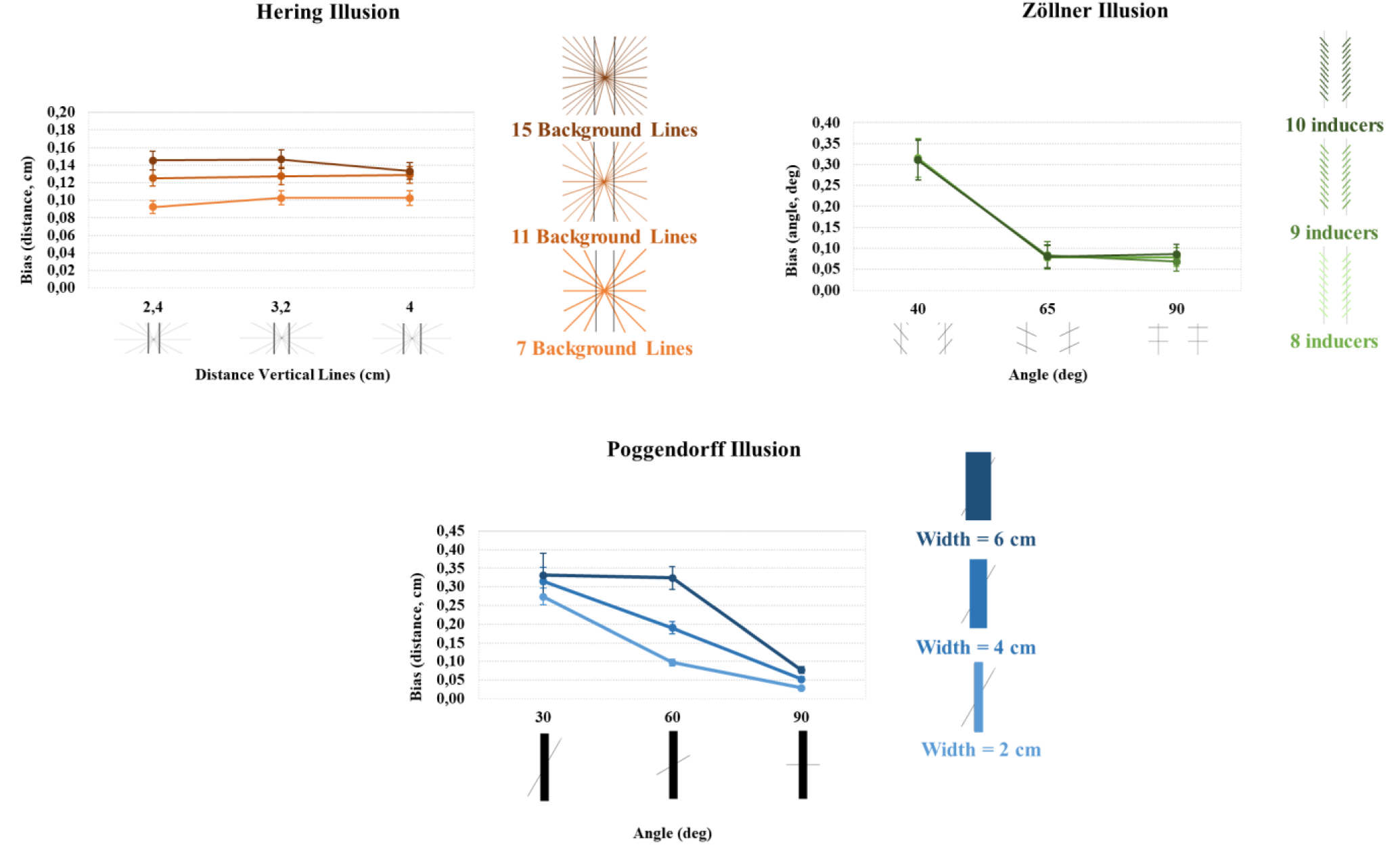
Behavioral Results. **Hering illusion:** Mean sizes of illusion for the three different number of background lines are shown as a unction of the three distances between the two vertical lines. Error bars: standard error of the mean. **Zöllner illusion:** Mean biases (in bsolute value) for the three different number of inducers are shown as a function of the three levels of angles. Error bars: standard rrors. **Poggendorff illusion:** Mean sizes of illusion (in absolute value) for the three different widths of the central surface are shown as function of the three levels of angles. Error bars: standard errors.

#### Hering illusion

There was a main effect of the number of radial lines (F (1.54, 44.63)=121.997, p<0.001, η^2^_p_=0.808), indicating that the participants’ bias increased as the number of radial lines increased: the more the radial lines, the larger the bias. 7 lines (average bias = 0.0992), 11 lines (average bias = 0,1272), 15 lines (average bias = 0,1417). No significant effect of the distance between the vertical lines was observed (F (1.29, 37.42)=0.8, p=0.4). However, there was a significant interaction between the number of radial lines and the distance between the vertical lines (F (2.9, 84.17)=5.79, p<0.01, η^2^_p_=0.166). Bias was significantly above 0 in all conditions (Hering: t(29) >= 12.30, p < .001). Inspection of the results suggests that whereas for the medium number of lines (11), the distance between the vertical lines has no effect on the bias, for both the 7 lines and the 15 lines conditions distance affects the bias but in opposite ways. Post-hoc one-way ANOVAs that examined the effect of distance for each line condition, separately, confirmed these observations. Specifically, no effect of distance was found for the 11 lines condition. In contrast, a significant effect of distance was found for both the 7 lines condition (F(1.34, 38.91)=5.46, p=0.01, η^2^ _p_ =0.16) and the 15 lines condition (F(1.49, 43.2)=3.45, p=0.05, η^2^ _p_ =0.106). In the case of the 7 lines condition, participants’ bias was significantly smaller for the 2.4cm distance condition (bias=0.0923) compared to both 3.2cm (bias =0.1025) and 4cm (bias =0.1027) (p=0.01) (Table 2). Finally, in the case of the 15 lines condition, participants’ bias was found to be significantly smaller for the largest distance condition (bias=0.1332) compared to the medium distance condition (bias=0.1464) (p<0.05) (Table 2). The bias for the smallest distance condition was very similar to the medium distance one (bias=0.1455) (Table 2).

**Table 2.** Behavioral and computational results for Hering illusion. Average bias and standard errors are shown per stimulus

| Radial lines | Vertical lines distance | Averaged behavioral bias (distance, cm) | Standard error of behavioral bias | Predicted computational bias (distance, pixels) |
| --- | --- | --- | --- | --- |
| 15 lines | 2.4 cm | 0.1455 | 0.0107 | 2.41 |
|  | 3.2 cm | 0.1464 | 0.0107 | 2.51 |
|  | 4 cm | 0.1332 | 0.0096 | 3.01 |
| 11 lines | 2.4 cm | 0.1253 | 0.0088 | 2.83 |
|  | 3.2 cm | 0.1273 | 0.0100 | 2.76 |
|  | 4 cm | 0.1288 | 0.0093 | 3.19 |
| 7 lines | 2.4 cm | 0.0923 | 0.0073 | 3.08 |
|  | 3.2 cm | 0.1025 | 0.0081 | 2.70 |
|  | 4 cm | 0.1027 | 0.0083 | 2.85 |

#### Zöllner illusion

There was a significant main effect of inducer angle on the measured bias (F (1.18, 34.29)=105.309, p<0.001, η^2^_p_=0.784), indicating that the illusion size is larger when the inducers’ angle is 40° (average bias=−0.312°) compared to both 65° (average bias=0.081°) and 90° (average bias=0.078°) (see for comparison Table 3). Neither the effect of the number of inducers (F (2, 58)=0.384, p=0.683, η^2^_p_=0.013) nor the interaction between the two factors (F (4, 116)= 0.486, p=0.746, η^2^_p_=0.013) were significant. Bias was significantly above 0 when lines inclination was 90° (t(29) >= 3.02, p < .005) and 65° (t(29) >= 2.56, p < .016; stats for 8,9,10 lines). In the illusory conditions – i.e. when lines inclination was 40° – bias was below 0 (t(29) <= −6.48, p < .001), Table 3.

**Table 3.** Behavioral and computational results for Zöllner illusion. Average bias and standard errors are shown per stimulus.

| Number inducers | Inducers' angle | Averaged behavioral bias (deg) | Standard error of behavioral bias | Predicted bias computational (deg) |
| --- | --- | --- | --- | --- |
| 10 inducers | 40° | -0.3108 | 0.048 | 0.3918 |
|  | 65° | 0.0806 | 0.027 | 0.0953 |
|  | 90° | 0.0858 | 0.024 | 0.0000 |
| 9 inducers | 40° | -0.3110 | 0.047 | 0.2899 |
|  | 65° | 0.0838 | 0.033 | 0.0957 |
|  | 90° | 0.0686 | 0.023 | 0.0000 |
| 8 inducers | 40° | -0.3156 | 0.046 | 0.3940 |
|  | 65° | 0.0779 | 0.028 | 0.1942 |
|  | 90° | 0.0785 | 0.022 | 0.0000 |

#### Poggendorff illusion

There was a significant main effect of the rectangle’s width on the measured bias (F (1.5, 33.3)=13.319, p=0.001, η^2^_p_=0.315). Participants’ bias increased as the rectangle’s width increased: 2cm (average bias = −0.114cm), 4cm (average bias = −0.151cm), 6cm (average bias=−0.193cm). A significant effect of the angle of the line segments was also observed (F (1.4, 40.87)=67.167, p<0.001, η^2^_p_=0.698). The smaller the angle, the larger the bias: 30° (average bias =−0.307cm), 60° (average bias=− 0.204cm), and 90° (average bias=0.053cm). Furthermore, the interaction between these two factors was also significant (F(1.98,57.56)=19.193,p<0.001, η^2^_p_=0.398). Bias was significantly above 0 in control conditions (i.e., 90°; t(29) >= 6.67, p < .001) and significantly below 0 in the illusory conditions (i.e. 30° and 60°, t(29) <= −5.52, p < .001). Inspection of the results suggested that the width of the rectangle affected participants’ bias in the 60° and 90° angles condition. Post-hoc one-way ANOVAs that examined the effect of rectangle’s width for each angle condition separately demonstrated that whereas the rectangle’s width had no effect on the participants’ bias in the 30° angle, in both 60° and 90°, rectangle’s width significantly affected the bias (F(1.13, 32.82)=63.34, p<0.001, η^2^_p_=0.686 and F(1.13, 32.85)=68, p<0.001, η^2^_p_=0.701, respectively). In both 60° and 90° conditions, participants’ bias increased as the rectangle’s width increased (Table 4). In 60°: 2cm (bias = −0.098cm), 4cm (bias = −0.190cm), 6cm (bias=−0.324cm), all significantly different from each other (p<0.001); and in 90°: 2cm (bias=0.029cm), 4cm (bias=0.052cm), 6cm (bias=0.076cm), all significantly different from each other, p<0.001, Table 4.

**Table 4.** Behavioral and computational results for Poggendorff illusion. Average bias and standard errors are shown per stimulus

| Central width | Crossing line angle | Averaged behavioral bias (distance, cm) | Standard error of behavioral bias | Predicted bias (distance, pixels) |
| --- | --- | --- | --- | --- |
| 6 cm | $30^\circ$ | -0.3310 | 0.0599 | 60 |
| | $60^\circ$ | -0.3242 | 0.0305 | 12 |
| | $90^\circ$ | 0.0765 | 0.0091 | 0 |
| 4 cm | $30^\circ$ | -0.3153 | 0.0376 | 36 |
| | $60^\circ$ | -0.1904 | 0.0164 | 6 |
| | $90^\circ$ | 0.0524 | 0.0064 | 0 |
| 2 cm | $30^\circ$ | -0.2742 | 0.0231 | 18 |
| | $60^\circ$ | -0.0976 | 0.0088 | 6 |
| | $90^\circ$ | 0.0288 | 0.0043 | 0 |

### Computational model results

#### Hering illusion

Predicted biases for this illusion are shown in Table 2. An example of a constructed percept is shown in Figure 6. Predicted biases were calculated as detailed in the *Methods section*. For 15 radial lines, the bias increases as the distance between the two vertical lines does: 2.4cm (bias = 2.41 px), 3.2cm (bias = 2.51 px), 4cm (bias = 3.01 px). In the 11 background lines condition, the computed bias is similar for the 2.4 cm and 3.2 cm conditions: 2.83 px and 2.76 px, respectively; for the 4 cm case the bias is 3.19 px. For 7 radial lines, the bias oscillates around the same values: 2.4cm (bias = 3.08 px), 3.2cm (bias = 2.70 px), 4 cm (bias = 2.85 px). The computed bias slightly increases as the number of radial lines passes from 7 to 11: 7 lines (average bias = 2.88 px), 11 lines (bias = 2.93 px). It decreases in the 15 lines example (bias = 2.64 px). The distance between the two vertical lines seems to play a role on the predicted bias in the computational approach when the distance passes from 3.2 to 4 cm: 2.4cm (average bias = 2.77 px), 3.2cm (average bias = 2.65 px), 4cm (average bias = 3.01 px).

**Figure 5.**
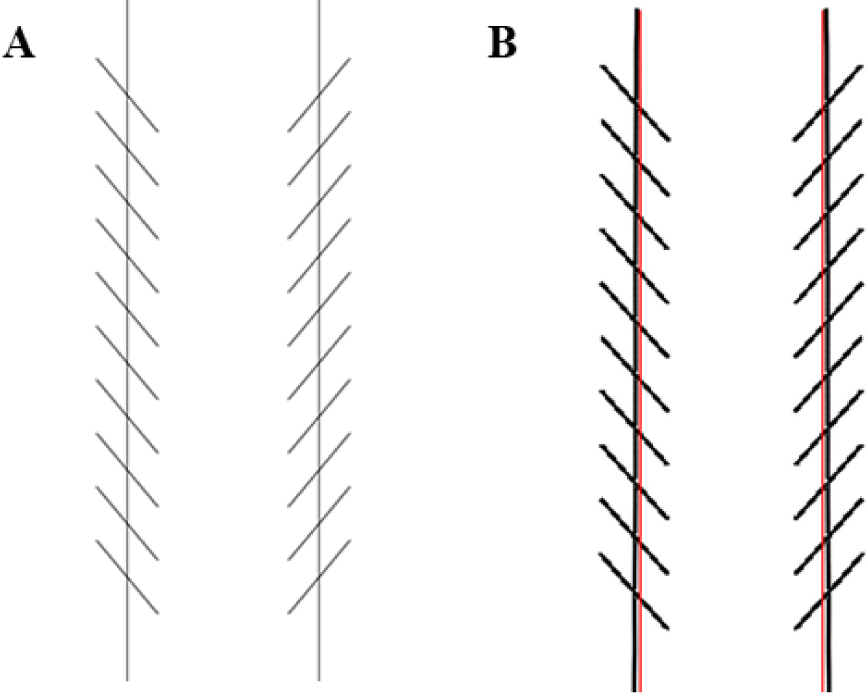
Zöllner stimuli: 10 inducers of 40°. **A.** Original stimulus. **B.** Constructed stimulus. In red, the two original vertical lines are shown

**Figure 6.**
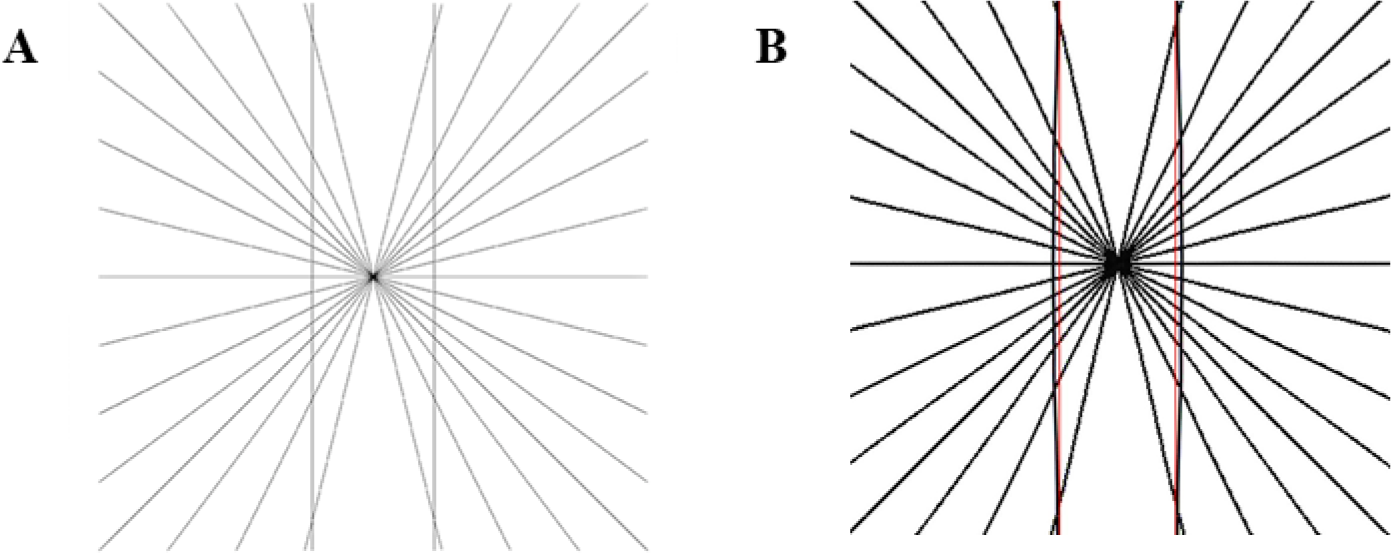
Hering stimuli: 15 background lines and 4 cm of distance between the vertical lines. **A.** Original stimulus. **B.** Constructed stimulus. In red, the two original vertical lines are shown, obtained with c = 0.02.

#### Zöllner illusion

Computed biases for this illusion are shown in Table 3. An example of a constructed percept is shown in Figure 5. For 10 inducers, the bias decreases from the 40° condition to the 65° condition: 40° (bias=0.3918°), 65° (bias = 0.0953°); and it goes to zero in the 90° condition. For 9 inducers, the bias decreases from the 40° condition to the 65° condition. In the case of angle of the inducers equal to 40°, the computed bias is 0.2899°, for 65° is 0.0957°. It goes to 0° when the angle is 90°. For 8 inducers, the bias is 0.3940° when the angle of inducers is 40° and it decreases to 0.1942° when the angles are 65°, to 0° when the angle is 90°.

There is a clear relation between the computed bias and the angle of the inducers, since the illusory bias increases when the inducers’ angle decreases from 40° (average bias =0.36°) to both 65° (average bias = 0.1284°) and 90° (average bias =0.0000°). However, there is no clear relation between the bias and the number of inducers on the predicted bias: 10 inducers (average bias = 0.16°), 9 inducers (average bias = 0.13°), 8 inducers (average bias = 0.1961°).

#### Poggendorff illusion

Modelled biases for this illusion are shown in Table 4. An example of a constructed percept is shown in Figure 7. For a central surface width of 6 cm, the bias decreases as the angle increases: 30° (bias= 60 pix), 60° (bias = 12 px), 90° (bias = 0 px). For the 4 cm width condition, the same happens: 30° (bias = 36 px), 60° (bias = 6 px), 90° (bias = 0 px). And as well for the 2 cm width: 30° (bias = 18 px), 60° (bias = 6 px), 90° (bias = 0 px). Comparing the predicted biases based on the central surface width, the bias increases as the width of the rectangle increases: 2 cm (average bias = 8 px), 4 cm (average bias = 14 px), 6 cm (average bias = 24 px). Whereas, when the angle of the crossing line is considered, the bias decreases when the angle increases: 30° (average bias = 38 px), 60° (average bias = 8 px), 90° (average bias = 0 px).

**Figure 7.**
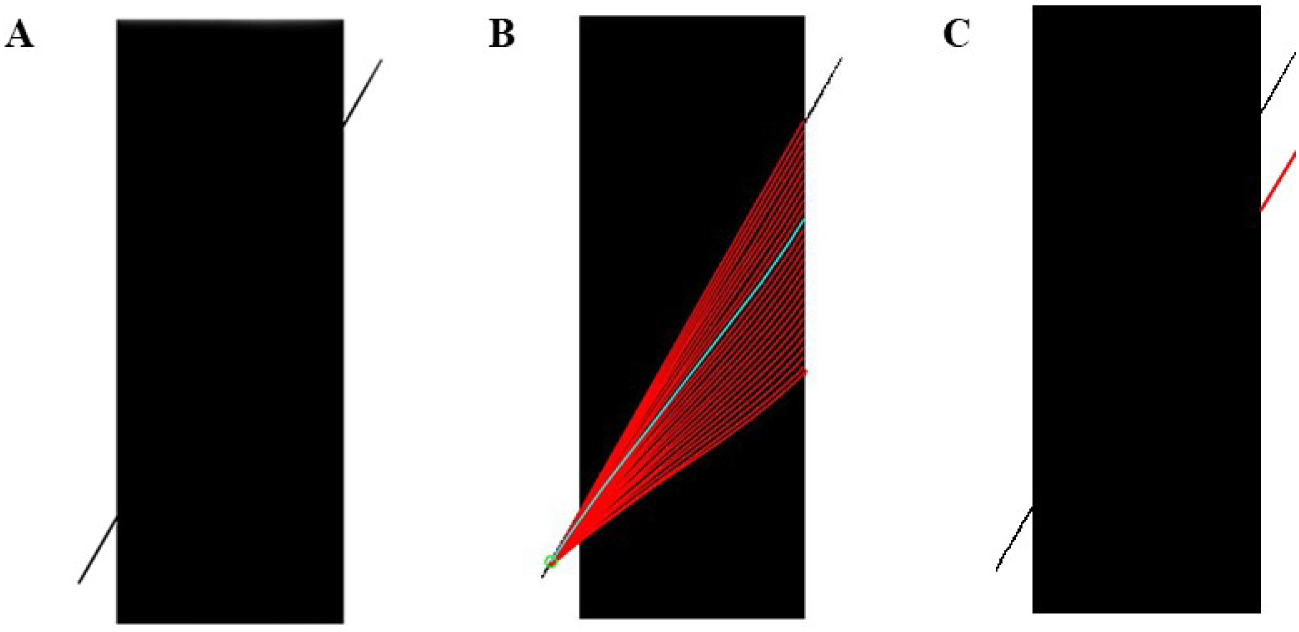
Poggendorff stimuli: 30° for the crossing line angle and 6 cm of central surface width. **A.** Original stimulus. **B.** Computed geodesics. The perceptual curve is shown in cyan. **C.** Constructed stimulus. In red, the computed percept

### Comparison behavioral and computational results

The comparison between the results of the two approaches: behavioral and computational results are shown in Figure 8, Figure 9 and Figure 10, listed in Table 5, Table 6 and Table 7, and detailed by geometrical optical illusion (Hering, Zollner and Poggendorff, respectively. B stands for behavioral curve, C for computational and D for difference curve in the figures). A comprehensive 3D visualization of the differences between behavioral and computational biases across illusion is reported in Figure 11. The comparison refers to the difference between the tuned computational model, as detailed in the corresponding Methods section, and the behavioral results. An example of not-fitted computational curve is given for the Hering illusion (c = 0.017) in Figure 8, where it is clearly depicted how the behavioral values serve as benchmarks to run multiple simulations of the computational model until an optimal fitting is reached (c = 0.037). In the Hering illusion, the difference curves are flat and differ overall by less than 0.50% (which represents about 30% of the maximum perceived bias), with the best fit achieved in the 11 lines case. Both of the computational models used here to mimic the perceptual behavior rely on the simulation of neurophysiological processes such as single cells responses and long-range connectivity in the visual areas. These models deal with different GOIs in a similar fashion to how the human visual system responds, making the differences across GOIs look similar for human behavior and computational modeling.

**Table 5.** Behavioral and computational results for Hering illusion. The difference between the results of both approaches is shown in the last column. The biases are shown per stimulus.

| Radial Lines | Vertical Lines Distance | Behavioral bias (percentage) | Computed bias (percentage) | Difference (percentage) |
| --- | --- | --- | --- | --- |
| 15 lines | 2.4 cm | 1.82 | 1.24 | 0.58 |
|  | 3.2 cm | 1.83 | 1.29 | 0.54 |
|  | 4 cm | 1.67 | 1.55 | 0.11 |
| 11 lines | 2.4 cm | 1.57 | 1.46 | 0.11 |
|  | 3.2 cm | 1.59 | 1.43 | 0.16 |
|  | 4 cm | 1.61 | 1.64 | 0.03 |
| 7 lines | 2.4 cm | 1.15 | 1.59 | 0.44 |
|  | 3.2 cm | 1.28 | 1.39 | 0.11 |
|  | 4 cm | 1.28 | 1.64 | 0.36 |

**Table 6.** Behavioral and computational results for Zöllner illusion (in absolute values). The difference between the results of both approaches is shown in the last column. The biases are shown per stimulus.

| Number Inducers | Inducers' Angle | Behavioral bias (deg) | Computed bias (deg) | Difference (deg - abs) |
| --- | --- | --- | --- | --- |
| 10 inducers | 40° | 0.3108 | 0.3918 | 0.081 |
|  | 65° | 0.0806 | 0.0953 | 0.0147 |
|  | 90° | 0.0858 | 0.0000 | 0.0858 |
| 9 inducers | 40° | 0.3110 | 0.2899 | 0.0211 |
|  | 65° | 0.0838 | 0.0957 | 0.0120 |
|  | 90° | 0.0686 | 0.0000 | 0.0686 |
| 8 inducers | 40° | 0.3156 | 0.3940 | 0.0784 |
|  | 65° | 0.0779 | 0.1942 | 0.1163 |
|  | 90° | 0.0785 | 0.0000 | 0.0785 |

**Table 7.** Behavioral and computational results for Poggendorff illusion (in absolute values). The difference between the results of both approaches is shown in the last column. The biases are shown per stimulus.

| Central Width | Crossing Line Angle | Behavioral bias (percentage) | Computed bias (percentage) | Difference (percentage) |
| --- | --- | --- | --- | --- |
| 6 cm | 30° | 2.32% | 16% | 13.78% |
|  | 60° | 2.27% | 3.21% | 0.94% |
|  | 90° | 0.54% | 0% | 0.54% |
| 4 cm | 30° | 2.21% | 9.63% | 7.42% |
|  | 60° | 1.33% | 1.60% | 0.27% |
|  | 90° | 0.37% | 0% | 0.37% |
| 2 cm | 30° | 1.92% | 4.81% | 2.90% |
|  | 60° | 0.68% | 1.60% | 0.92% |
|  | 90° | 0.20% | 0% | 0.20% |

**Figure 8.**
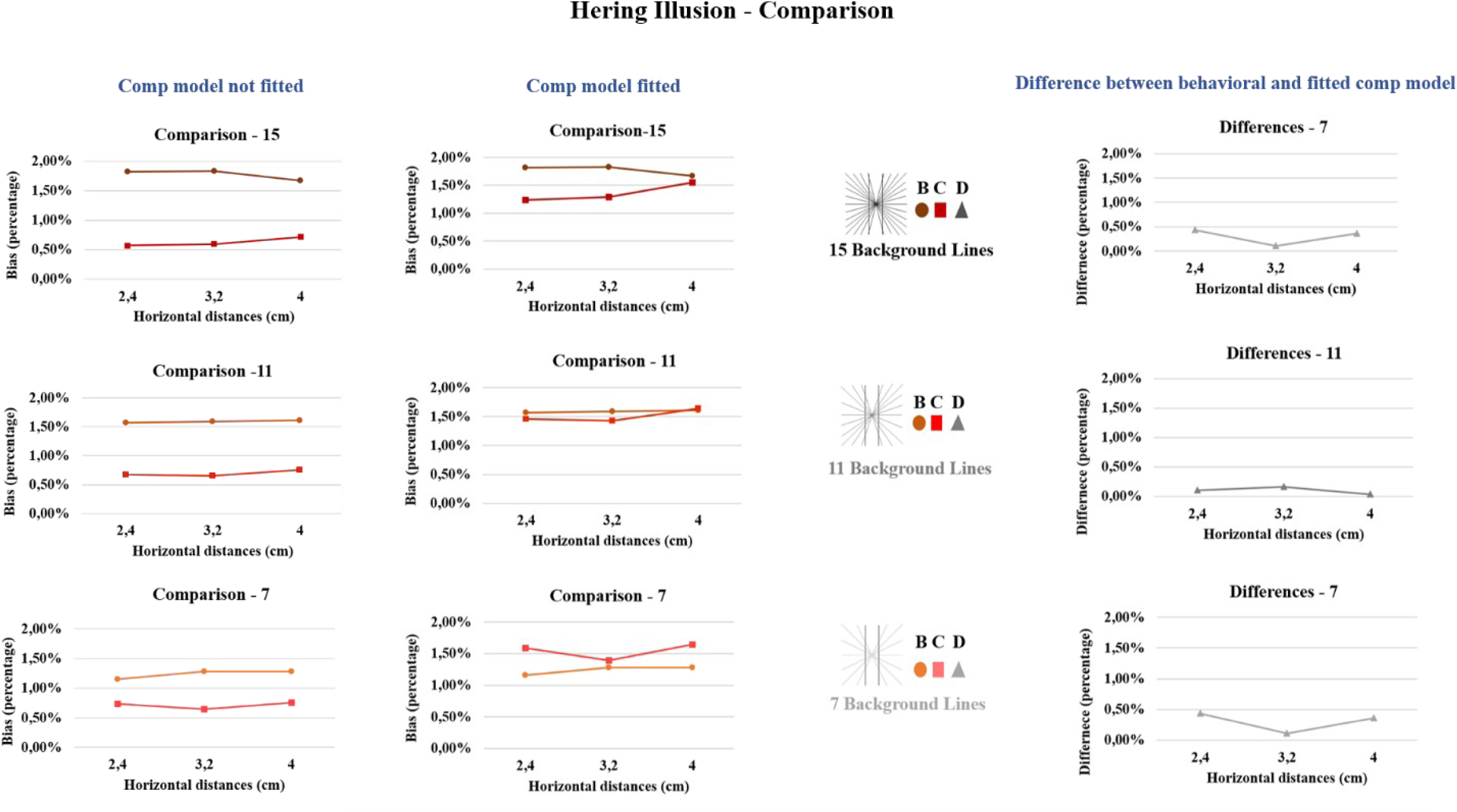
Comparison computational and behavioral results for the Hering illusion. **Left block:** Comparison of the behavioral and computational biases, in the left column the computational model does not fit the behavioral results (c = 0.017) while in the right column of the block it does (c = 0.037). In the computational cases − values for the predicted bias for the different number of background lines are shown as a function of the three distances between the two vertical lines. In the behavioral − mean sizes of illusion for the three different number of background lines are shown as a function of the three distances between the two vertical lines. **Right:** Differences between the behavioral and computational biases fitting the behavioral values. Values for the differences between the observed bias and the predicted bias for the different number of background lines are shown as a function of the three distances between the two vertical lines.

**Figure 9.**
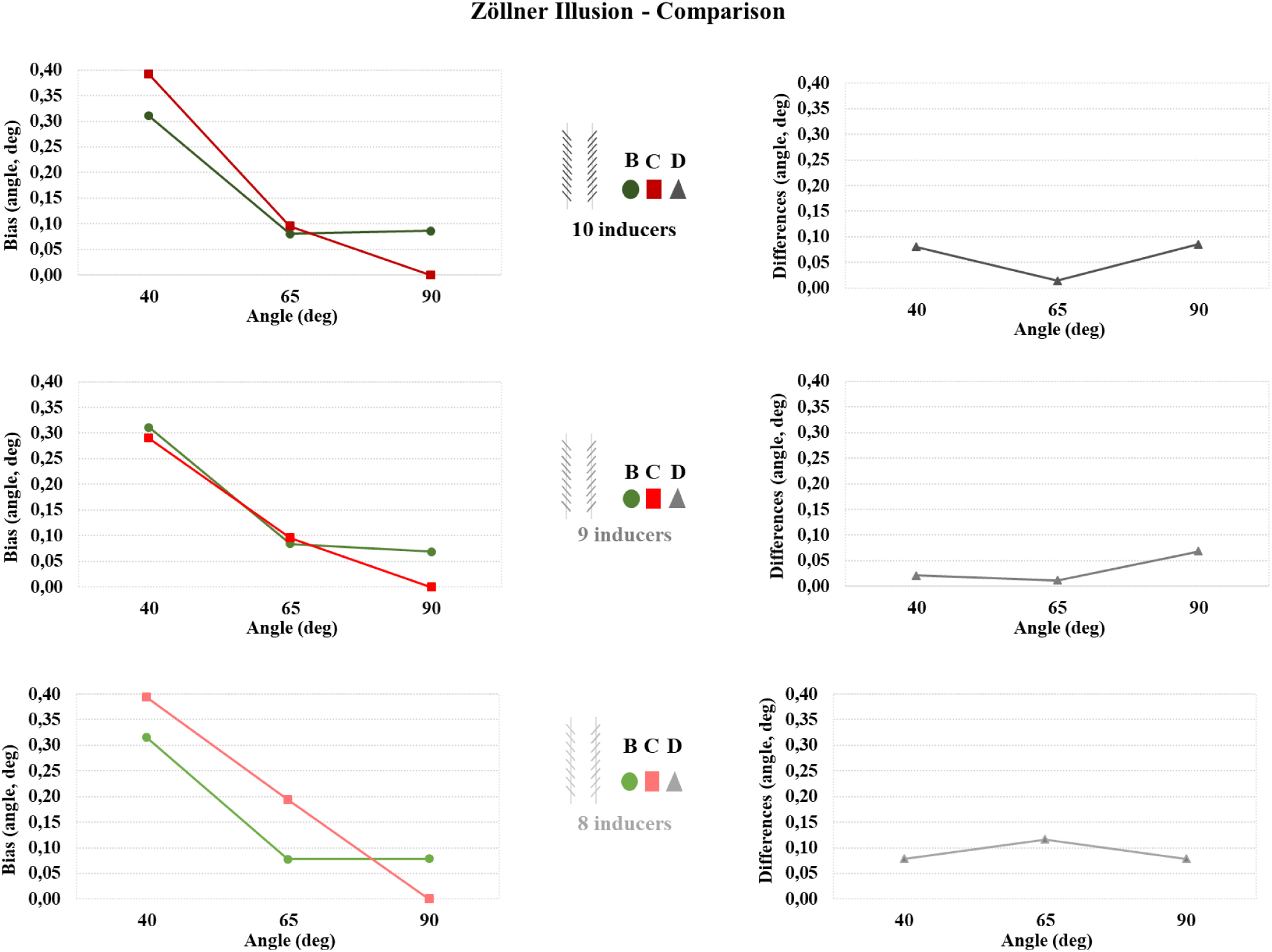
Comparison computational and behavioral results for the Zöllner illusion. **Left:** Comparison of the behavioral and computational biases. Computational − Absolute values for the predicted bias for the three different number of inducers are shown as a function of the three levels of angles. Behavioral − Mean biases (in absolute value) for the three different number of inducers are shown as a function of the three levels of angles. **Right:** Differences between the behavioral and computational biases’ values. Values for the differences between the observed bias and the predicted bias for the different number of inducers are shown as a function of the three different angles for the inducers.

**Figure 10.**
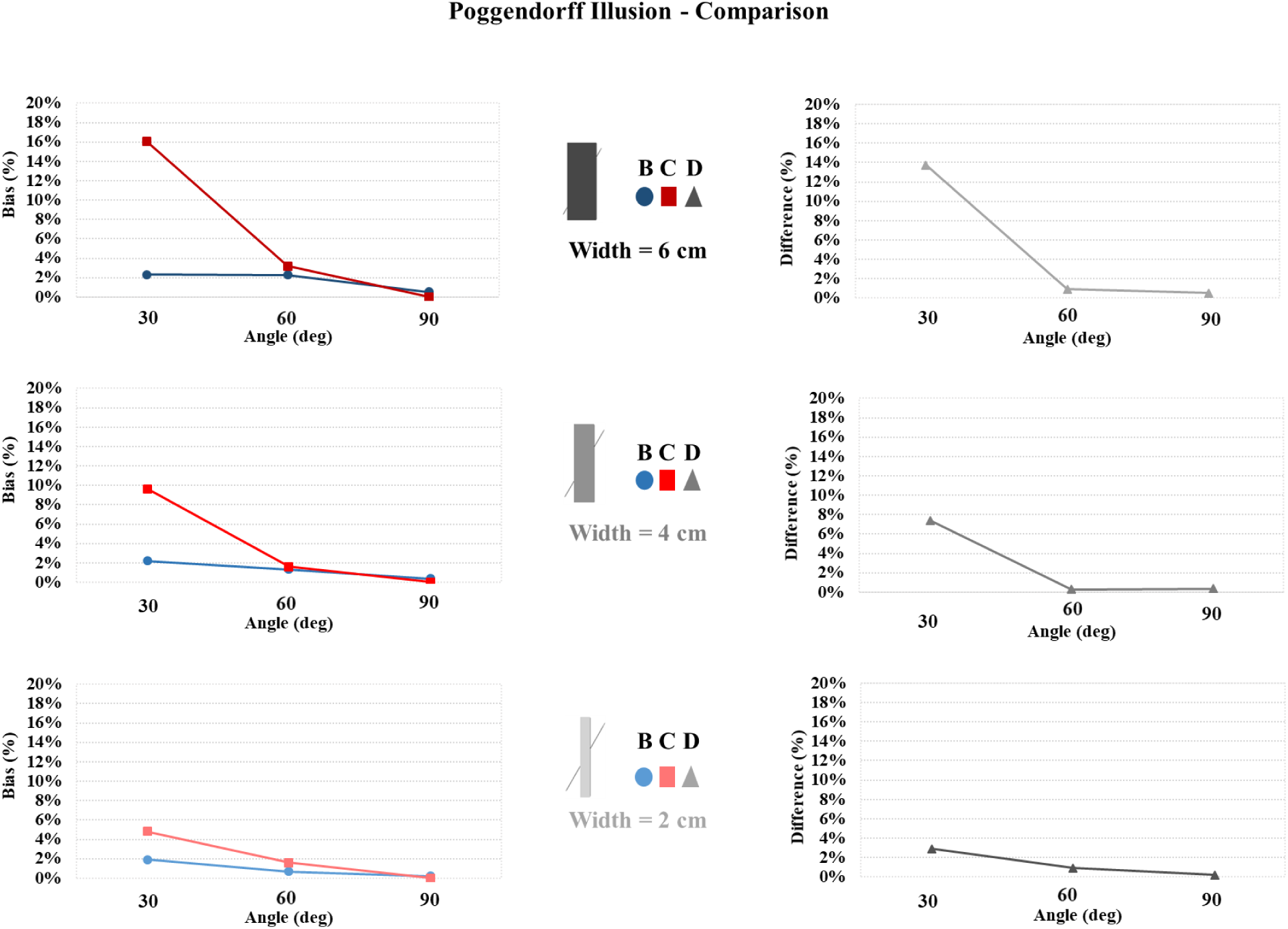
Comparison computational and behavioral results for the Poggendorff illusion. **Left:** Computational − Absolute values for the predicter bias for the three different widths of the central surface are shown as a function of the three levels of angles. Behavioral − Mean sizes of illusion (in absolute value) for the three different widths of the central surface are shown as a function of the three levels of angles. **Right:** Differences between the behavioral and computational biases’ values. Values for the differences between the observed bias and the predicted bias for the different widths of the central surface are shown as a function of the three different angles for the crossing line.

**Figure 11.**
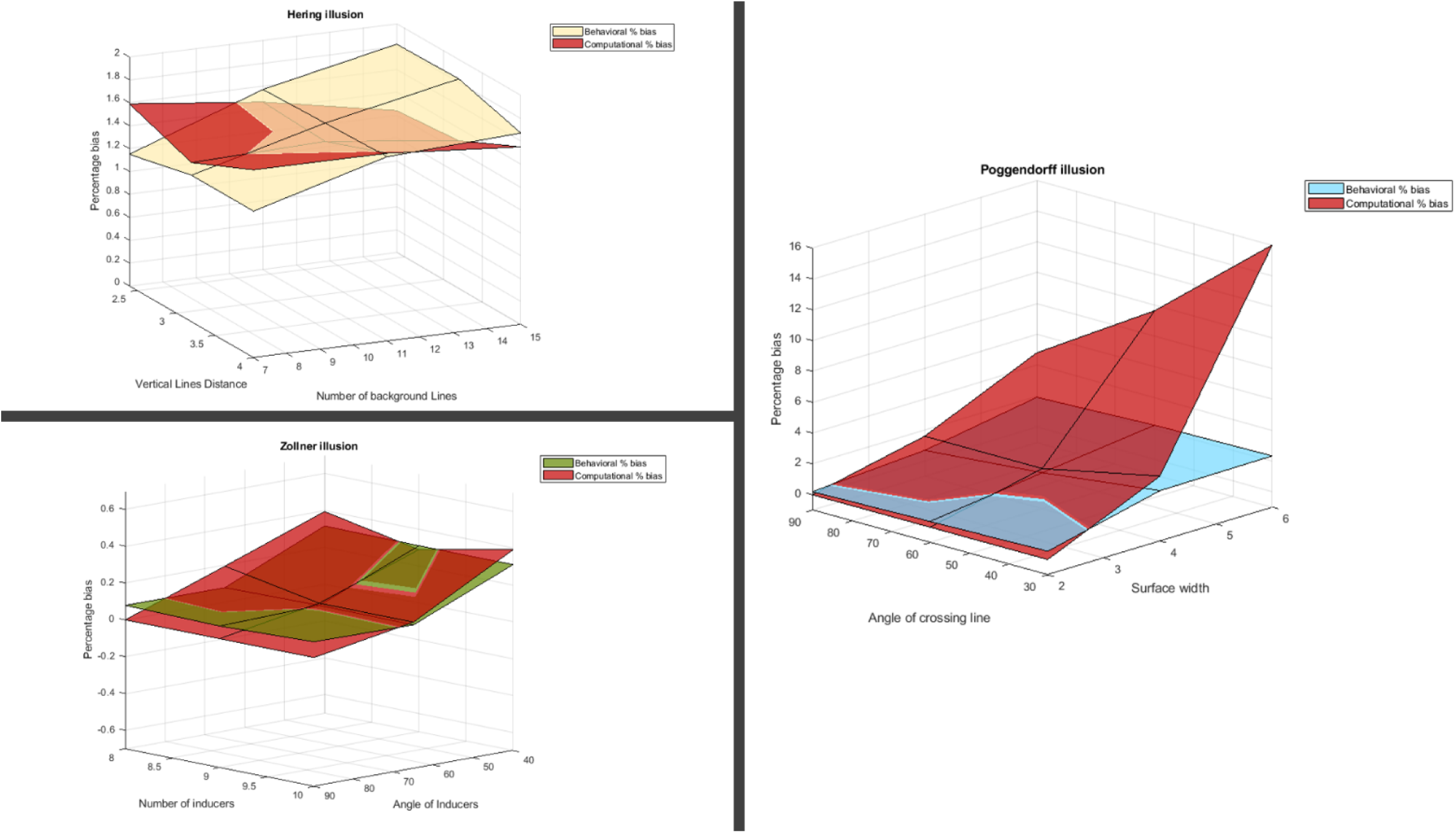
3D visualization of the differences between behavioral and computational bias for each of the three illusions.

## Discussion and conclusions

Three different GOIs have been studied in the present work: the Hering, the Zöllner and the Poggendorff illusions from psychophysical and computational standpoints. The recorded behavioral biases for the three illusions served as a basis for tuning the parameters of the computational model in order to obtain computational curves replicating numerically the behavioral biases. Our results provide a roadmap whereby computational modelling, informed by human psychophysics, can reveal likely mechanistic underpinnings of perception.

To the best of our knowledge, this was the first time these three illusions were behaviorally studied together and that a standardized definition of bias was quantified, with up-to-date experimental methods. The determinants of the biases in these three GOIs share an angular component given by the intersection of the foreground line (or surface) with radial lines, segments (inducers) or one crossing line (Hering: number of radial lines and distance between the vertical lines; Zöllner: the number and angle of inducers; Poggendorff: width of the central surface and the angle of the crossing line). This angular component seems to reflect the intracortical connectivity of hypercolumns: neurons belonging to the same hypecolumn – and spiking simultaneously – might be a first determinant of misperception (bias). Second, the distance between lines (or the width of the surface) is depicted by the global integration of local selected features, therefore being more representative of the way through which long-range connectivity represents visual stimuli.

With this in mind, our results indicate that for the Hering illusion, the number of radial lines does influence the perception; more radial lines resulted in larger bias, in agreement with Holt-Hasen, 1961 (Holt-Hansen, 1961). A similar effect was observed for the Zöllner illusion: the inducers’ angle had an effect on the perceived bias, which increases as the intersect angle decreases, in agreement with observations in Oyama, 1973 (Oyama, 1975). On the other hand, no significant main effect of distance between the two vertical lines was reported in the Hering, only an interaction between the two variables indicating that the distance was affecting participants’ performance in the minimum and maximum radial lines conditions but not the medium one. If we consider the perceptually more complex phenomenon of the Poggendorff illusion, its determinants, i.e. the angle of the crossing line and the width of the central surface, both had an influence on the perception of the illusion, in accordance with previous studies (Jones-Buxton & Wall, 2001; Weintraub & Krantz, 1971). In fact, the participants’ bias increased as the rectangle’s width increased, while the smaller the angle, the larger the bias. This seems to suggest that the presence of a rectangle surface constitutes a much stronger determinant for the formation of long-range connections than two straight lines (Hering illusion). Also, an interaction between the two factors (width and angle) at a behavioral level was observed. It is also worthwhile to observe that in the behavioral results of the three illusions, the observed bias is always significantly different from zero. This would indicate that participants do not have a sharp “non-illusory” effect while tested. Instead, while observers do indeed have the perceptual experience of what is not an illusion, there is nonetheless a slight bias reported. We did not find a relationship between the sizes of the biases across the three different illusions: this suggests that the described mechanisms (intracortical and long-range connectivity) contribute to a different extent to these illusions.

The behavioral results served as a basis for tuning the computational models, i.e. iteratively exploring the set of parameters representing the intra-cortical (*σ*; *γ*; 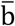 – Gabor filters parameters) and long-range connectivity (c; ξ) within the model to produce a computational bias matching the recorded behavior. Computational models have the unique value of enabling the *in-silico* exploration of the boundaries of the neurophysiological mechanisms modelled without the need of running an *in vivo* experiment to do so. Within the same illusions, all predicted (computational) biases have been computed by fixing the size of receptive profiles and the constant *c*, related to *α*^−1^ of Formula 8. The size of receptive fields both mediates for local interaction between simple cells belonging to the same hypercolumn, together with playing the role of a subsequent level of visual cortices (V2), known to have larger receptive fields. A fine tuning of the parameters detailed above helped to compromise between the contribution of intra-cortical connectivity and long-range connectivity: the need for large-size (*σ*) receptive fields has been identified by means of the model, pointing out the likely contribution of higher-level visual processes such as those in V2 and perhaps elsewhere (Murray & Herrmann, 2013b). The strength of neuro-geometrical models give a clear and elegant framework to explain the cellular organization and construction of visual percepts, which can be easily applied to other illusions, such as the Kanizsa triangle (Citti & Sarti, 2006) or basic perceptual phenomena (line completion) (Favali, Citti, & Sarti, 2017).

The computational models replicated well the Zollner behavioral results as well as specific conditions of both the Poggendorff and Hering illusions. Undoubtedly, the current neuro-geometrical models despite the fact that they approximate the neurophysiological time features of long-range connectivity, i.e. the speed at which such connections take place in the hypercolumnar structure of V1 and V2, they cannot accomplish a perfect correspondence with neurophysiology. However, despite its limitations, the computational model provides insight into the relevant neurological processes likely underlying these illusions. Its effectiveness in predicting “spatial-related” phenomena such as the Zollner illusion, where any effect of distance between the vertical lines seems to play a central role, indicates that the main neural mechanism involved is this illusion is the intra-cortical connectivity of V1, which is thought to transpire on a relatively fast timescale. On the other hand, it is also worthwhile to consider those conditions that were poorly fitted in the Poggendorff illusion. Here, the effect of the width of the rectangle seemed to produce a larger bias compared to the one obtained from the behavioral results. This suggests that further investigating the temporal component of perception of these stimuli, (i.e. feedback mechanisms from higher-level visual cortices into primary visual cortex), might improve our understanding of how visual neuronal processes create such images. Our computational and behavioral results could therefore be complemented by brain mapping and neuroimaging studies with high temporal resolution, such as EEG (Biasiucci, Franceschiello, & Murray, 2019).

A limitation of the computational modelling implemented here is that it was not possible to generate variability in performance that is similar to the inter-individual variability observed in human participants. This could be artificially generated via Monte-Carlo simulations able to automatically generate a variance around the Gabor filters’ parameters or the long-range connectivity parameters. However, this would presuppose that the parameters affecting Gabor filters or long-range connectivity are the only factors causing the illusory effects.

To conclude, the present study combining a carefully designed behavioral and computational paradigms highlights the likely neurophysiological mechanisms e.g. intra-cortical connectivity and long-range connectivity in V1/V2 implicated in the Hering, Zollner and the Poggendorff illusions and provides a roadmap for future studies using psychophysical data to tune computational models.

## Acknowledgments

Financial support for this work has been provided by the Fondation Asile des Aveugles (grant #232933 to M.M.M.), a grantor advised by Carigest SA (#232920 to M.M.M.), as well as the Swiss National Science Foundation (grants #169206 to M.M.M.). We thank Dr. Jonas Richiardi for helpful comments on an earlier version of this manuscript.

## Authors contributions

C.R., M.M.M. and B.F. conceptualized the problem. N.N. contributed to the development of the behavioral experiment including implementation of software for rendering the illusions and collecting behavioral data. C.R. and B.F. tested the protocol and collected the data. L.R. provided the mathematical formulation for the bias in the Hering illusion, as described in the appendix, and the formal analysis of the phenomena. C.R. performed the behavioral analysis. A.H.A. performed the computational analysis, elaborated the graphs and performed the behavioral versus computational comparison, under C.R. and B.F.’s supervisions. C.R. M.M.M. and B.F. drafted the manuscript, and all authors contributed to internal review.

## Materials and correspondence

Correspondence and requests for materials should be addressed to B.F. or C.R.

## Appendix

### A) Mathematical expression for the curves manipulated manually by the participant

In the Hering illusion the stimulus consists of a pair of vertical straight lines, on top of a collection of straight lines passing through a central point. The observer perceives the straight vertical lines as slightly curved away from the central point in the horizontal direction. Our goal is to identify an equation for the perceived curve that can be manipulated within an experimental design.

This illusion has some clear symmetries, namely flipping across the vertical axis or across the horizontal axis does not change anything. Therefore we can reduce our consideration to the top right quadrant and then extend by reflection.

In this quadrant we will consider curves leaving a fixed point on the top edge and reaching the bottom edge at a variable point and being orthogonal to both edges at the intersection point. The first point and the tangent directions are fixed, but the position of the second point is what makes the curve deviate from the neutral configuration (i.e. when the curves are vertical straight lines). The subjects can choose the position of the second endpoint until it matches to what they perceive as straight, given an initial offset (see Methods section).

Since we are imposing three fixed conditions (one control point and two tangent directions), we need a family of curves with at least four degrees of freedom; three degrees are fixed by these three conditions, and the subject has one degree of freedom to play with. The easiest such family is given by the algebraic curves defined by polynomials of degree at most three and real coefficients (this family is a real vector space of dimension 4; lower degree would not give us enough flexibility; higher degree would result in extra degrees of freedom).

In order to set up our computations, let us choose the vertical axis as the x-axis, oriented upwards, and the horizontal axis as the y-axis, oriented to the right. For simplicity, we are going to align the x-axis along the right vertical straight line in the neutral configuration. In this way the curves we care about will intersect the y-axis in points of the form (0,B) for some *B* ∈ ℝ, *with B =* 0 corresponding to the neutral configuration, and they will intersect the x-axis at the endpoint (C,0), where C is half of the edge length of the original picture, equal to 45% of the total screen height where the experiment was performed. While the parameter C is fixed, the value of B is precisely what the subject manipulates.

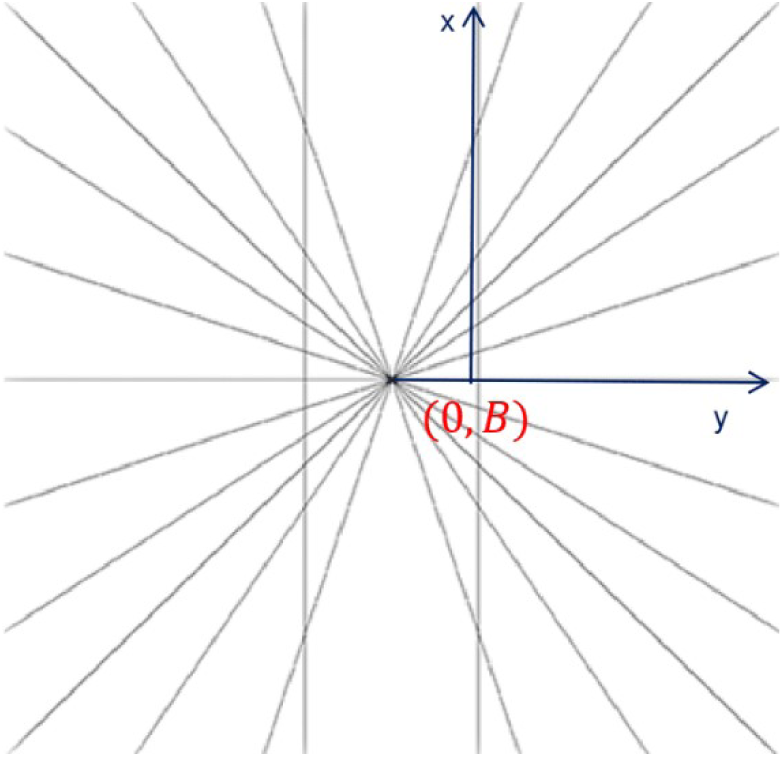

The equation for such a polynomial of degree 3 in these coordinates is given by

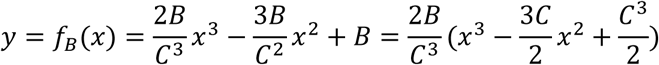

Once again, here C is a fixed quantity, while B is allowed to vary, describing the 1-parameter family of curves we need. The choice of B is equivalent to some geometrically meaningful quantities, which can be computed from the first and second derivatives.

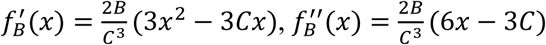

For instance, notice that the coordinates of the unique point of flex of the above curves are

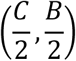

and the tangent line at this point has equation

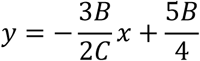

which means it has slope 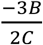. To get another geometric parameter, notice that the curvature at the endpoints can be computed in terms of the second derivative at those points, which is

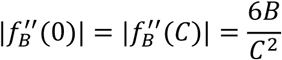

It should be remarked that in our setting *B* << *C*, so that both the slope and the curvature are essentially zero, hence not useful from the numerical point of view, even though they are a good theoretical measure of how far these curves are from the straight lines. To have a numerically useful parameter we stick therefore to B, namely the distance in cm (or bias) along the whole paper; we emphasize once again that the choice of B is equivalent to these geometric features.

